# Pentamycin biosynthesis in Philippine *Streptomyces* sp. S816: Cytochrome P450-catalysed installation of the C-14 hydroxyl group

**DOI:** 10.1101/604736

**Authors:** Shanshan Zhou, Lijiang Song, Joleen Masschelein, Felaine A.M. Sumang, Irene A. Papa, Teofila O. Zulaybar, Aileen B. Custodio, Daniel Zabala, Edwin P. Alcantara, Emmanuel L.C. de los Santos, Gregory L. Challis

**Affiliations:** Department of Chemistry, University of Warwick, Coventry CV4 7AL, United Kingdom; National Institute of Molecular Biology and Biotechnology (BIOTECH), University of the Philippines Los Baños, Los Baños, Laguna 4031, Philippines; Warwick Integrative Synthetic Biology Centre, University of Warwick, Coventry CV4 7AL, United Kingdom; Department of Biochemistry and Molecular Biology, Monash University, Clayton, Victoria 3800, Australia

## Abstract

Pentamycin is a polyene antibiotic, registered in Switzerland for the treatment of vaginal candidiasis, trichomoniasis and mixed infections. Chemical instability has hindered its wide-spread application and development as a drug. Here we report the identification of *Streptomyces* sp. S816, isolated from Philippine mangrove soil, as a pentamycin producer. Genome sequence analysis identified the putative pentamycin biosynthetic gene cluster, which shows a high degree of similarity to the gene cluster responsible for filipin III biosynthesis. The *ptnJ* gene, which is absent from the filipin III biosynthetic gene cluster, was shown to encode a cytochrome P450 capable of converting filipin III to pentamycin. This confirms that the cluster directs pentamycin biosynthesis, paving the way for biosynthetic engineering approaches to the production of pentamycin analogues. Several other *Streptomyces* genomes were found to contain *ptnJ* orthologues clustered with genes encoding polyketide synthases that appear to have similar architectures to those responsible for the assembly of filipin III and pentamycin, suggesting pentamycin production may be common in *Streptomyces* species.

---

Polyene antibiotics are a subgroup of macrolides that are particularly active against fungi. They target sterols in fungal cell membranes, destabilizing them, which leads to eventual cell death.^1^ Polyenes have polyunsaturated macrolactone core structures that are assembled by type I modular polyketide synthase (PKS) assembly lines.^1^ Diversification of the core structures results from post-PKS tailoring reactions, such as O-glycosylation and C-hydroxylation.^1^

Filipin III^2^ (**1**) and pentamycin^3^ (**2**) are closely-related polyenes containing 28-membered macrolides that differ only by a hydroxyl group at C-14 in the latter (Figure 1). Filipin III displays similar affinity for membranes containing ergosterol (the primary sterol in fungal membranes) and cholesterol (the primary sterol in mammalian membranes) ^1^, making it unsuitable for use as a drug. However, it is used clinically in the diagnosis of type C Niemann-Pick disease, which results in the accumulation of cholesterol in the lysosome, due to its high intrinsic fluorescence and cholesterol-binding ability.^4^ Pentamycin is active against *Candida albicans*, *Trichomonas vaginalis* and several pathogenic bacteria.^5^ It is registered in Switzerland for the treatment of vaginal candidiasis, trichomoniasis and mixed infections,^6^ and has also been shown to increase the efficacy of the anti-cancer drug bleomycin.^7^ However, chemical instability has hindered the wide-spread application of pentamycin and its further development as a drug.^8^

**Figure 1.**
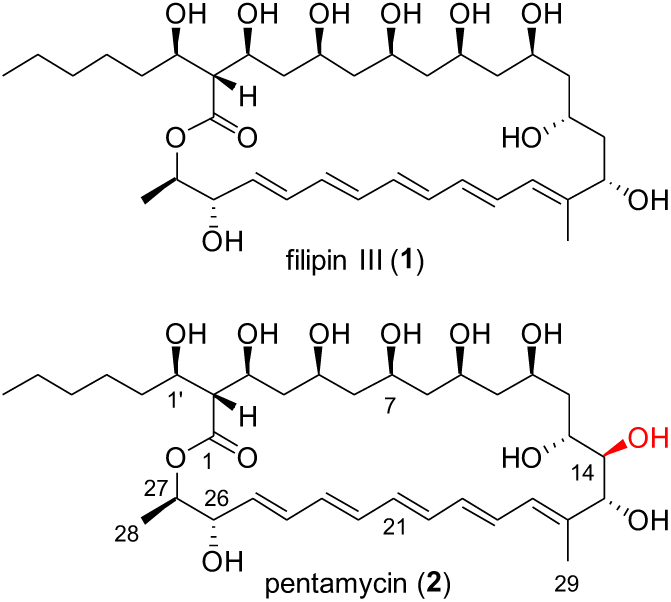
The structures of filipin III (**1**) and pentamycin (**2**). The only difference is the addition of a hydroxyl group (highlighted in red) to C-14 in the latter.

Filipin III was originally isolated from *Streptomyces filipinensis* DSM40112, but has since been reported to be produced by other *Streptomyces* strains, including *Streptomyces avermitilis* MA-4680.^9^ The filipin biosynthetic gene cluster (*pte*) in *S. avermitilis* encodes a type I modular PKS containing 14 modules distributed across 5 subunits and 8 additional proteins that govern regulation of its expression and tailoring of the polyketide skeleton.^10^ The *pte* cluster contains two genes (*pteC* and *pteD*) encoding cytochromes P450 (CYPs) that have been shown to catalyze hydroxylation of C-26 and C-1’, respectively. The molecular basis for the regio- and stereospecificity of these transformations has been elucidated by X-ray crystallography.^11^

Here we report that *Streptomyces* sp. S816, previously isolated from Philippine mangrove soil,^12^ produces pentamycin. Sequencing and analysis of this strain’s genome identified the putative pentamycin biosynthetic gene cluster (*ptn*), which shows a high degree of similarity to the *pte* cluster. An additional CYP-encoding gene in the *ptn* cluster was hypothesised to be responsible for introduction of the C-14 hydroxyl group. This hypothesis was confirmed by showing that the CYP can convert filipin III to pentamycin. Identification of the *ptn* cluster will facilitate future efforts to produce more stable derivatives of pentamycin via engineering of its biosynthetic pathway.

## RESULTS AND DISCUSSION

### Isolation and initial characterisation of *Streptomyces* sp. S816

The isolation of *Streptomyces* sp. S816 from mangrove soil collected in Surigao del Sur in the Philippines and its morphological characterization have been reported previously.^12^ Morphological traits include septate, branched and granular aerial substrate mycelia with yellow pigmentation. Among 135 actinobacterial isolates from Philippine soils collected during the study, the S816 strain was found to be the most effective against methicillin-resistant *Staphylococcus sciuri* isolated from the milk of cows and goats suffering from mastitis. The degree of growth inhibition by extracts from this strain was higher than that caused by vancomycin. It was thus selected for further characterisation.

### Identification of pentamycin as a metabolite of *Streptomyces* sp. S816

Methanol extracts of *Streptomyces* sp. S816 cultures on Soy Flour Mannitol agar were analysed by UHPLC-ESI-Q-TOF-MS, revealing a compound with *m/z* = 671.4000, congruent with the molecular formula of pentamycin (C_35_H_58_O_12_; calculated *m/z* = 671.4001 for C_35_H_59_O_12_^+^; see supporting information). The UV-Vis absorbance spectrum of this compound had λ_max_ values at 324, 339 and 357 nm, identical to those reported in the literature for pentamycin (see supporting information).^13^ UHPLC-ESI-QTOF-MS comparisons with an authentic standard provided further confirmation that *Streptomyces* sp. 816 produces pentamycin (see supporting information).

### Genome sequencing of *Streptomyces* sp. S816 identifies the putative pentamycin biosynthetic gene cluster

Genomic DNA of *Streptomyces sp.* S816 was isolated and sequenced using the Illumina MiSeq platform via a combination of paired end and mate pair reads. SPaDES^14^ was used to assemble the two libraries into a high-quality draft genome sequence. The sequence was annotated using DFAST (see supporting information for further details) and the autoMLST^15^ pipeline was used to construct a phylogenetic tree from alignments of core genes in closely related genomes. This confirmed our previous taxonomic classification of the S816 strain as a member of the *Streptomyces* genus, closely related to *S. murinus* (Figure 2).

**Figure 2.**
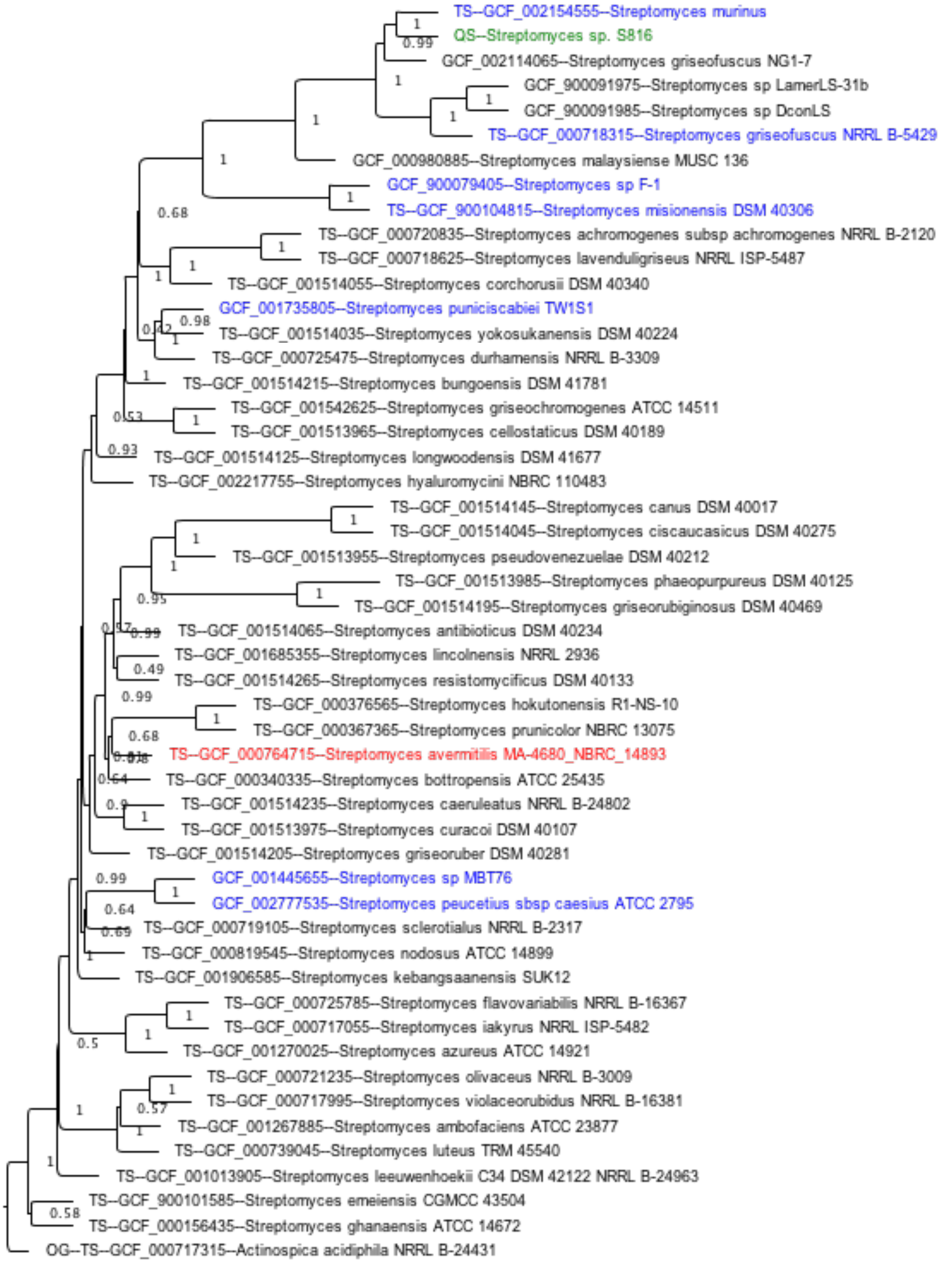
Phylogenetic comparison of *Streptomyces* sp. S816 with other *Streptomyces* species. The consensus phylogenetic tree was built from a multi-locus alignment of core genes in *Streptomyces* sp. S816 (green) with related species in the NCBI Assembly database, including the filipin III producer, *S. avermitilis* MA-4680 (red), and strains containing *ptnJ* orthologues (blue).

To assess the strain’s potential to produce specialised metabolites, the genome sequence was analysed using antismash,^16^ which identified thirty-four putative biosynthetic gene clusters. One of these showed a high degree of similarity to the *S. avermitilis* filipin III biosynthetic gene cluster (Figure 3 and Table1) and was hypothesised to direct pentamycin bioynthesis.^17^ The module and domain organisation of the type I modular PKS encoded by this cluster is congruent with the carbon skeleton of pentamycin (Figure 4).

**Figure 3.**
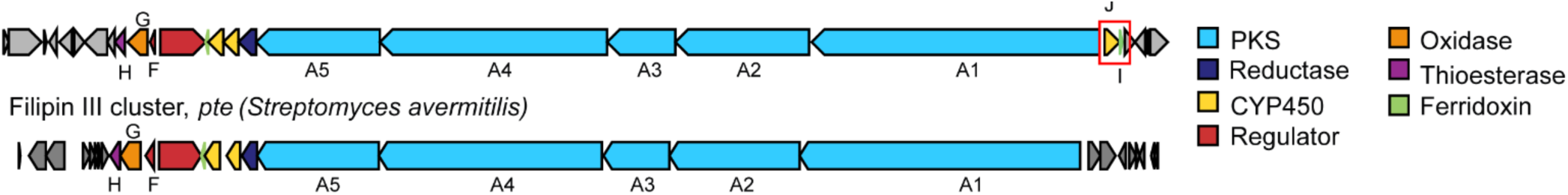
Comparison of the pentamycin and filipin III biosynthetic gene clusters in *Streptomyces* sp. 816 and *S. avermitilis* MA-4680, respectively. The two clusters show a high degree of synteny. Genes encoding a putative CYP and ferredoxin (highlighted by the red box) lie upstream of the PKS genes in the pentamycin cluster. These two genes are absent from the filipin III cluster.

**Table 1.** Proposed functions of genes in the *Streptomyces* sp. S816 pentamycin biosynthetic gene cluster and comparison with genes in the *S. avermitilis* MA-4680 filipin III biosynthetic gene cluster.

| *Streptomyces* sp. S816 |  |  | *S. avermitilis* MA-4680 |  |  |
| --- | --- | --- | --- | --- | --- |
| Proposed Function | Gene Name (Gene ID) | Amino Acids | Gene Name (Gene ID) | Amino Acids | % Identity |
| ferredoxin for PtnJ | *ptnI* (DV517_01430) | 64 | - | - | NA |
| CYP (hydroxylation of C-14) | *ptnJ* (DV517_01420) | 405 | - | - | NA |
| PKS | *ptnA1* (DV517_01410) | 7840 | *pteA1* (SAV419) | 7746 | 84 |
| PKS | *ptnA2* (DV517_01400) | 3613 | *pteA2* (SAV418) | 3564 | 86 |
| PKS | *ptnA3* (DV517_01390) | 1823 | *pteA3* (SAV417) | 1835 | 84 |
| PKS | *ptnA4* (DV517_01380) | 6153 | *pteA4* (SAV416) | 6160 | 85 |
| PKS | *ptnA5* (DV517_01370) | 3353 | *pteA5* (SAV415) | 3352 | 86 |
| crotonyl-CoA reductase/carboxylase | *ptnB* (DV517_01360) | 418 | *pteB* (SAV414) | 418 | 93 |
| CYP (hydroxylation of C-1') | *ptnC* (DV517_01350) | 399 | *pteC* (SAV413) | 399 | 92 |
| CYP (hydroxylation of C-26) | *ptnD* (DV517_01340) | 404 | *pteD* (SAV412) | 416 | 89 |
| ferredoxin for PtnC/PtnD | *ptnE* (DV517_01330) | 80 | *pteE* (SAV411) | 64 | 81 |
| SARP-LAL regulator | *ptnR* (DV517_01320) | 1188 | *pteR* (SAV410) | 1171 | 79 |
| PAS-LuxR regulator | *ptnF* (DV517_01310) | 192 | *pteF* (SAV409) | 232 | 88 |
| cholesterol oxidase | *ptnG* (DV517_01300) | 547 | *pteG* (SAV408) | 536 | 90 |
| thioesterase | *ptnH* (DV517_01290) | 255 | *pteH* (SAV407) | 255 | 82 |

**Figure 4.**
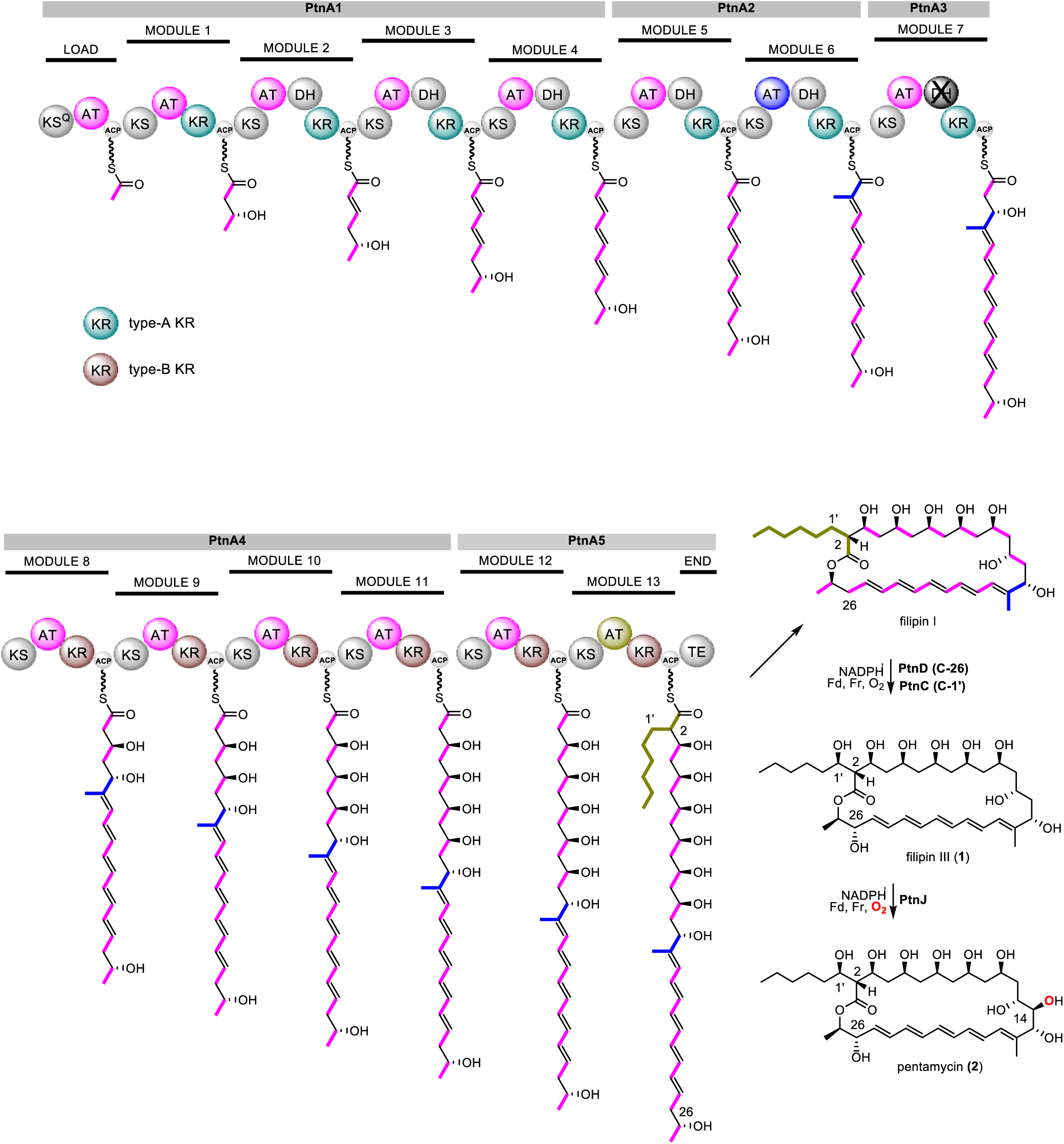
Proposed pathway for pentamycin biosynthesis in *Streptomyces* sp. S816. Domain abbreviations are as follows: KS ketosynthase, KS^Q^: malonyl-ACP decarboxylase; AT acyltransferase, ACP acyl carrier protein, DH dehydratase, KR ketoreductase, TE thioesterase. The C-1’ and C-26 hydroxylation reactions in filipin III biosynthesis can occur in either order.^4^ While PtnJ can hydroxylate C-14 of filipin III, the possibility that this reaction can also occur at earlier stages in the biosynthetic pathway cannot be excluded.

The putative pentamycin biosynthetic gene cluster contains two genes upstream of the first PKS gene, encoding a putative CYP (*ptnJ*) and ferredoxin (*ptnI*), which are absent from the filipin III biosynthetic gene cluster. We hypothesised that the CYP catalyses the hydroxylation of C-14 in pentamycin biosynthesis (Figure 4).

### PtnJ catalyses the conversion of filipin III to pentamycin

In order to directly examine the role played by PtnJ in pentamycin biosynthesis, we cloned and overexpressed the gene encoding this putative CYP in *Escherichia coli*. Recombinant PtnJ was produced as an N-terminal His_8_-tagged fusion protein, allowing straightforward purification by Nickel affinity chromatography (see supporting information). The UV-Vis spectrum of the purified protein had an absorbance maximum at 418 nm, which is typical for CYPs in their ferric resting state. This shifted to 450 nm in the ferrous C=O complex, further confirming the sequenced-based annotation of PtnJ as a CYP (see supporting information).

The purified protein was incubated with filipin III, NADPH and spinach ferredoxin (Fd) / ferredoxin reductase (Fr) for 3 h at 25 °C. UHPLC-ESI-Q-TOF-MS analyses showed that the major product of the reaction has the same retention time and *m/z* value as pentamycin (Figure 5). This compound was absent from control reactions containing the heat-denatured CYP, or lacking NADPH, Fd, or Fr (Figure 5). Taken together, these data show that PtnJ catalyzes hydroxylation of C-14 in filipin III to form pentamycin.

**Figure 5.**
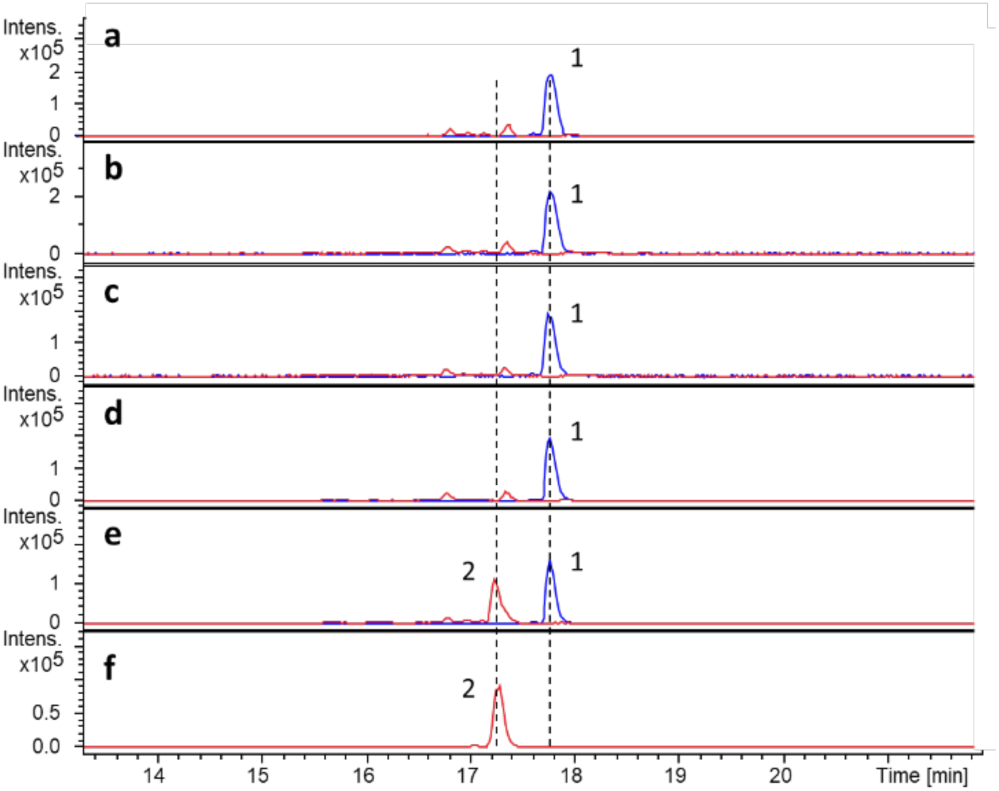
Extracted ion chromatograms at *m*/*z* = 693.3820 and 677.3871, corresponding to [M+Na]^+^ for pentamycin (red traces) and filipin III (blue traces), respectively, from UHPLC-ESI-Q-TOF-MS analyses of PtnJ-catalysed reactions. a: control -NADPH; b: control -Fd; c: control -Fr; d: control containing boiled PtnJ; e: enzymatic reaction; f: pentamycin standard.

### Pentamycin-like biosynthetic gene clusters are found in several other *Streptomyces* genomes

A BLASTP query of the NCBI non-redundant database using the sequence of PtnJ returned 7 orthologous (>85% identity) hits, all *Streptomyces* species (see supporting information). Analysis of the regions flanking the genes encoding these hits using antismash^16^ indicated that they are clustered with genes encoding type I modular PKSs with a high degree of similarity to those involved in pentamycin and filipin III biosynthesis. The species harbouring these gene clusters are phylogenetically diverse (Figure 2), suggesting pentamycin production may be common in the *Streptomyces* genus. The gene encoding the CYP responsible for hydroxylation of C-26 appears to be absent from the cluster in *Streptomyces peucetius* subsp. *caesius* ATCC27952 (see supporting information), indicating this strain produces a novel pentamycin analogue.

## CONCLUSION

Taken together, our data strongly support the hypothesis that PtnJ is a CYP that catalyzes the hydroxylation of C-14 in pentamycin biosynthesis. Identification of the pentamycin biosynthetic gene cluster paves the way for production of pentamycin analogues via biosynthetic engineering. Deletion of the genes encoding the CYPs responsible for the hydroxylation of C-26 (FilC) and C-1’ (FilD) in filipin biosynthesis leads to the production of filipin analogues with more potent antifungal activity.^4^ A similar strategy could be used to create pentamycin analogues lacking the hydroxyl groups at C-26 and/or C-1’. Indeed, such analogues may be produced naturally by *Streptomyces* species containing *ptnJ* orthologues clustered with genes encoding PKSs that appear to assemble the filipin III/pentamycin carbon skeleton. The pentamycin PKS could also be engineered to produce derivatives with modified carbon skeletons, which may possess greater chemical stability and thus be better suited to clinical application.

## ASSOCIATED CONTENT

Materials; growth and extraction of *Streptomyces* sp. S816 for metabolite analyses; genome sequencing assembly and annotation; overproduction, purification and UV-Vis spectroscopic analysis of PtnJ; analysis of PtnJ-catalysed hydroxylation on filipin III; bioinformatics identification and comparison of gene clusters containing PtnJ orthologues.

The *Streptomyces sp.* S816 genome sequence described in this paper has been deposited at DDBJ/ENA/GenBank under the accession QQVZ00000000. The version described in this paper is version QQVZ01000000.

## Author Contributions

G.L.C., E.L.C.dlS. and E.P.A. designed and coordinated the project. E.L.C.dlS. performed the genome assembly and bioinformatics analysis. J.M. designed the *ptnJ* expression vector and S.Z. purified and characterised PtnJ. L.S. identified pentamycin as a metabolite of *Streptomyces* sp. S816. F.A.S., D.Z., I.A.P, T.O.Z., A.B.C, and E.P.A., isolated *Streptomyces* sp. S816, grew cultures for metabolite profiling, and isolated genomic DNA. E.L.C.dlS., G.L.C., S.Z. and L.S. wrote the manuscript.

## Notes

The authors declare no competing financial interests.

## ACKNOWLEDGMENTS

This project was funded by the British Council through a Newton Institutional Links Award (Grant Ref. 261846416). The Bruker MaXis Impact UHPLC-ESI-Q-TOF-MS instrument used in this research was purchased with a grant from the BBSRC (BB/K002341/1 to G.L.C.). E.L.C.d.l.S. is a Research Career Development Fellow in the Warwick Integrative Synthetic Biology Centre, which is supported by a grant from the BBSRC and EPSRC (BB/M017982/1).

---

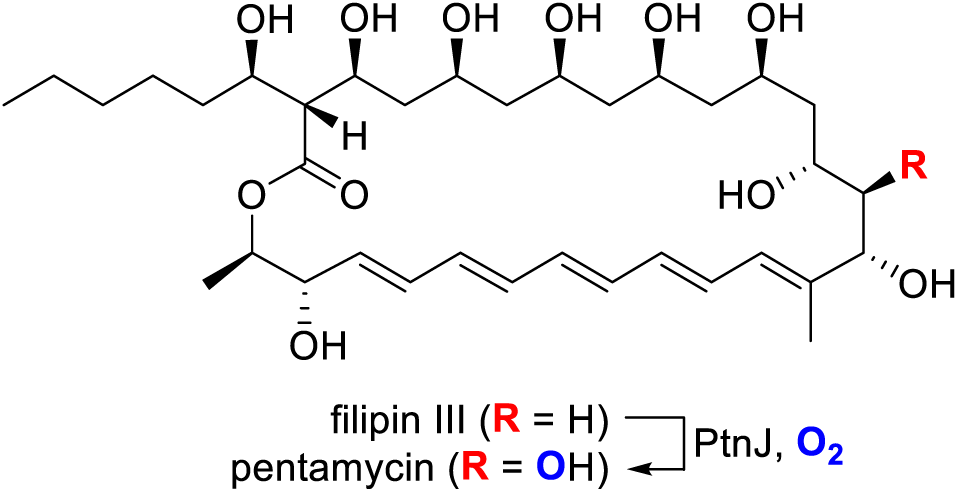

