## Supplemental Information for "Pentamycin biosynthesis in Philippine *Streptomyces* sp. S816: Cytochrome P450-catalysed installation of the C-14 hydroxyl group"

### Supporting information

#### Materials

Partially purified ferredoxin (1.0-3.0 mg/mL solution in 0.15 M Trizma buffer, pH 7.5 and NaCl) and ferredoxin reductase (8.5 units made to 0.5 mL in Tris 25 mM, pH 8, 0.1 M NaCl, 20% glycerol) from *Spinacia oleracea* were purchased from Sigma and were used without further purification.

#### Analysis of pentamycin production by *Streptomyces* sp. S816

A Soy Flour Mannitol agar[1] plate was inoculated with spores of *Streptomyces* sp. S816 and grown for 5 days at 30°C. The agar was chopped into small pieces and extracted with 3 x 25 mL of MeOH. The combined extracts were concentrated to dryness under reduced pressure and the residue was dissolved in 1 mL of MeCN/$H_2O$ (1:1 v/v). After filtration through a 0.2 μm filter, the resulting solution was analysed using a Dionex Ultimate 3000 RS UHPLC instrument equipped with ZORBAX Eclipse Plus C18 column (2.1 × 100 mm, 1.8 μm) coupled to a Bruker MaXis Impact mass spectrometer [ESI in positive ion mode; full scan 50-2500 *m/z*; end plate offset, -500 V; capillary, -4500 V; nebulizer gas ($N_2$), 1.4 bar; dry gas ($N_2$), 8 L/min; dry temperature, 200 °C]. The column was eluted with solvent A (0.1% formic acid in water) and solvent B (0.1% formic acid in acetonitrile) using the following elution profile at a flow rate of 0.2 mL/min: 0-5 min, 5% B; 5-20 min, linear gradient from 5% B to 100% B; 20-25 min, 100% B; 25-28 min, linear gradient from 100% B to 5% B; 28-34 min, 5% B.. The mass spectrometer was calibrated with 10 mM sodium formate at the beginning of each run. Pentamycin in the extracts had the same retention time and mass / UV spectra as an authentic pentamycin standard (>95% purity, AG Scientific) (**Figures S1-S3**).

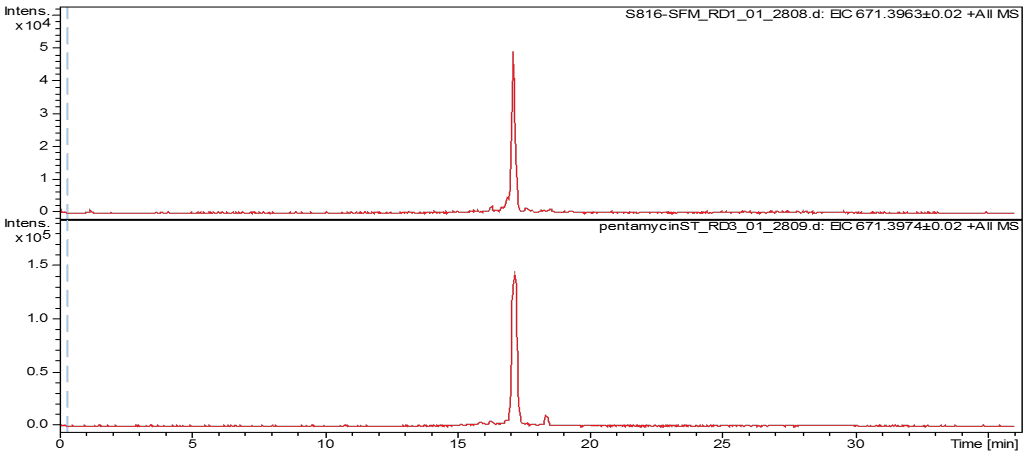


**Figure S1:** Comparison of pentamycin in extracts of *Streptomyces* sp.S816 (top chromatogram) with the authentic standard of pentamycin (bottom chromatogram).

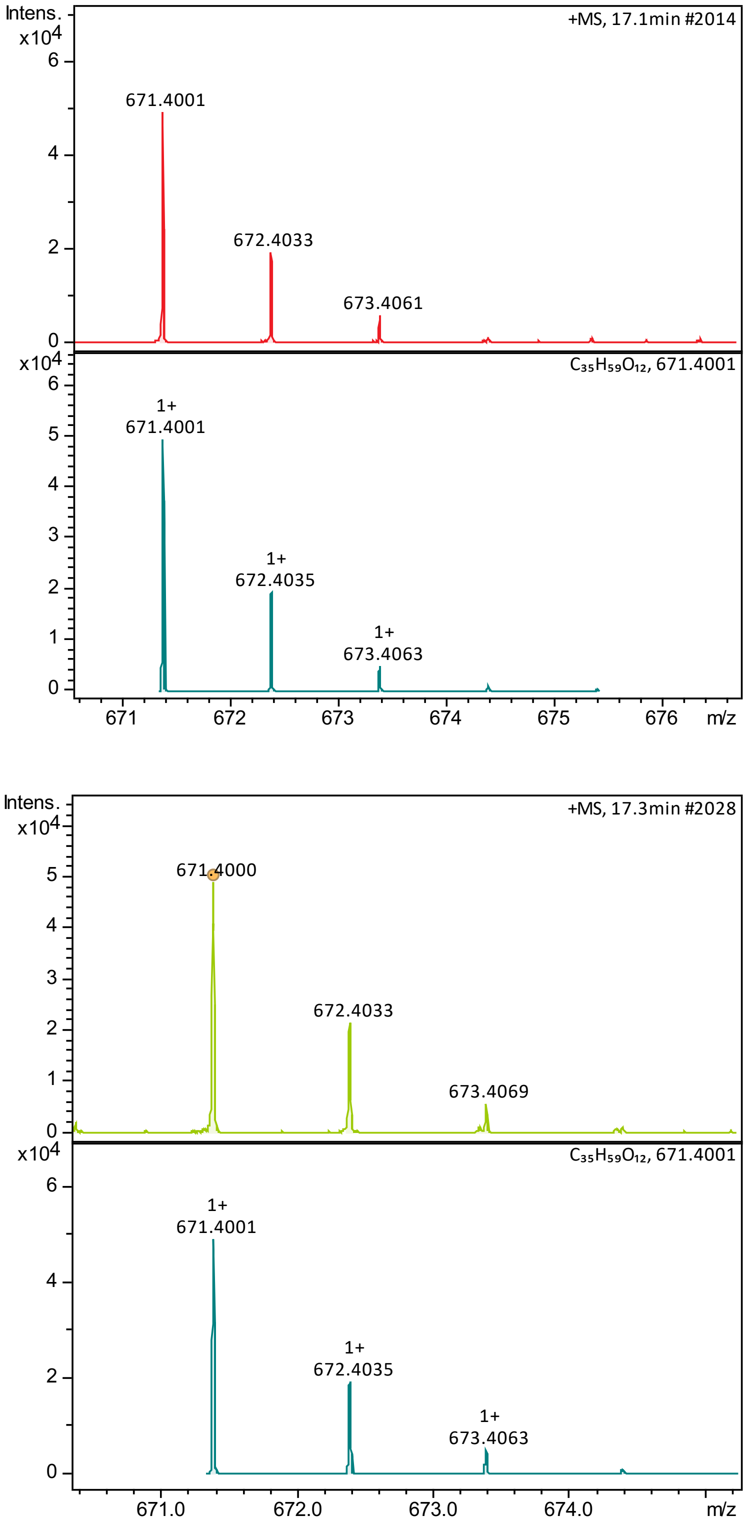


**Figure S2:** Comparison of measured and simulated mass spectra for pentamycin in extracts of *Streptomyces* sp. S816 (top) and the authentic standard of pentamycin (bottom).

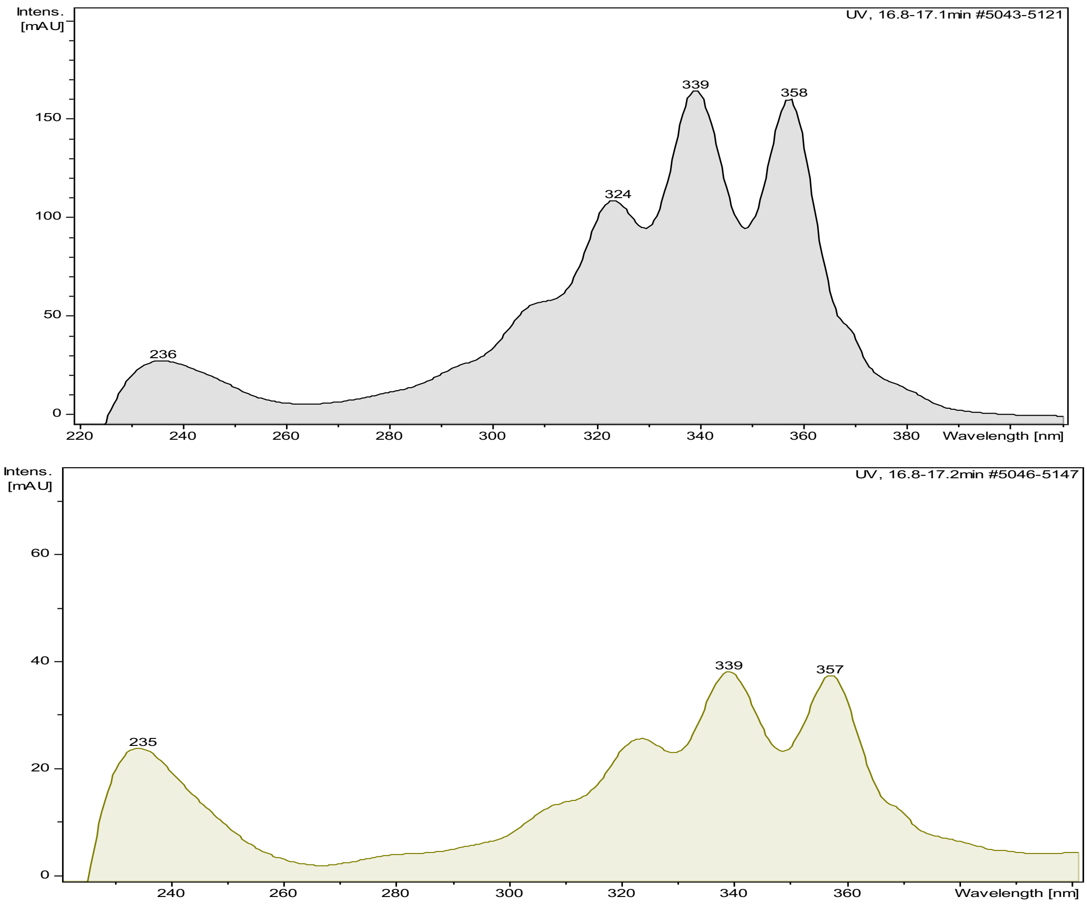


**Figure S3:** Comparison of UV-Vis spectra for pentamycin in extracts of *Streptomyces* sp. S816 (bottom) and the authentic standard of pentamycin (top).

#### Genome Assembly and Bioinformatics

A loopful of *Streptomyces* sp. S816 cells grown for 5 days at 30°C on tryptic soy agar was used to inoculate 10 ml of tryptic soy broth and the resulting culture was incubated for two days at 30°C and 200 rpm. The cells were pelleted by centrifugation at 12,000 rpm and 20°C and genomic DNA was extracted using a Qiagen Puregene kit from approximately 50-100 mg (wet weight) of pelleted cells according to the manufacturer's instructions. The extracted DNA was stored in DNA hydration solution (Qiagen) at -20°C.

The *Streptomyces* sp. S816 genomic DNA was sequenced by the DNA Sequencing Facility in the Department of Biochemistry at the University of Cambridge. A TruSeq PCR-free library, and a Nextera Mate Pair library were prepared and were sequenced in an Illumina MiSeq System using a 500 cycle v2 kit with 250 base pair paired end reads. Raw reads were trimmed using trim_galore[2] and assembled using SPaDes.[3] The assembled genome was annotated using DFAST[4]. antiSMASH[5] with the knownclusterblast option was run on the annotated genome, and the putative pentamycin biosynthetic gene cluster was identified on the basis of its similarity to the filipin III biosynthetic gene cluster.

Multilocus phylogenetic analysis was done in part using the autoMLST[6] pipeline. Briefly, autoMLST selects organisms to build a phylogenetic tree by estimating the average nucleotide identity of the query sequence to assemblies in the NCBI database. The core genome sequences of these organisms were extracted and aligned to the query and a consensus phylogenetic tree based on the individual alignments was built. In addition to the organisms selected by the autoMLST, genomes of organisms that contained BLAST hits with a high homology (> 75% identity) to *ptnJ* were added to the phylogenetic analysis.

**Overproduction and purification of PtnJ**

pET24a containing *ptnJ* with an in-frame $His_8$-encoding sequence fused to its 5' end was purchased from Epoch Life Science. *E. coli* BL21star (DE3) cells were transformed with the pET24a-*ptnJ* plasmid for expression of the gene. 15 µL of a glycerol stock of *E. coli* BL21star (DE3) cells carrying pET24a-*ptnJ* were used to inoculate 15 mL of LB medium containing 50 µg/mL of kanamycin and the resulting culture was incubated for 16 hours (37 °C, 180 rpm). 10 mL of this preculture was used to inoculate 1 L of LB medium (1% v/v) containing 50 µg/mL of kanamycin and the resulting culture was grown for 3-5 hours (37 °C, 180 rpm) until the optical density at 600 nm reached 0.6-0.8. Isopropyl-β-D-thiogalactopyranoside (final concentration 0.1 mM), $FeCl_3$ (final concentration 5 mg/L) and L-glutamic acid (final concentration 1 mM) were added, and the culture was grown for a further 16 hours (15 °C, 180 rpm).

The cells were harvested by centrifugation at 5000 rpm for 20 min and re-suspended in washing buffer (15 mL per litre culture) containing 20 mM Tris-HCl, pH 8, 100 mM NaCl, 20 mM imidazole and 10% v/v glycerol, and phenylmethanesuphonyl fluoride was added (final concentration 1 mM). The resuspended cells were lysed using a Constant Systems E1061 cell disrupter and centrifuged at 17,000 rpm for 30 min to separate insoluble material. The supernatant was passed through a 0.2 µm filter and loaded onto a pre-equilibrated 1 mL HisTrap HP nickel affinity column (GE Healthcare). Unbound proteins were removed by washing with 10 mL of the above buffer and $His_8$-PtnJ was eluted via successive washes with 3 mL of elution buffer (20 mM Tris-HCl, pH 8, 100 mM NaCl, 10% v/v glycerol) containing 50 mM, 100 mM, 200 mM and 300 mM imidazole. The fractions were analyzed by SDS-PAGE, and those containing the recombinant protein were collected and concentrated using a Vivaspin ultrafiltration column (Sartorius) with a 30 kDa cutoff. The purified protein, which was a yellow-orange color, was exchanged into an imidazole-free storage buffer (20 mM Tris-HCl, pH 8, 100 mM NaCl and 10% v/v glycerol) using a PD-10 column, and aliquots were flash frozen in liquid nitrogen and stored at -80 °C. SDS-PAGE confirmed the purity of the protein (**Figure S4**).

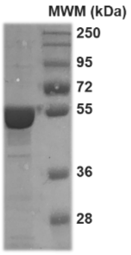


**Figure S4:** SDS-PAGE analysis of $His_8$-PtnJ after purification. Molecular weight marker (MWM): PageRuler Plus prestained protein ladder (ThermoFisher). The predicted molecular weight of $His_8$-PtnJ is 46,840 Da.

**UV-Vis spectroscopic analysis of purified $His_8$-PtnJ**

UV-Vis spectroscopy was carried out on a Perkin Elmer Lambda 35 UV/Vis spectrometer. PtnJ was confirmed as a cytochrome P450 by UV-Vis spectroscopic analysis of its ferrous C=O complex.[7,8,9]

Purified protein was diluted to 8 µM with Tris buffer (25 mM, pH 8) and the UV-Vis spectrum was measured before and after the addition of 5 µL of a 1 M solution of sodium dithionite (**Figure S5**). Carbon monoxide gas was bubbled through a separate sample of the protein solution for 2-3 min, sodium dithionite (5 µL, 1 M) was added and the UV-Vis spectrum was measured (**Figure S5**). The C=O difference spectrum was generated by subtracting the spectrum of the reduced protein in the absence of C=O from that of the reduced protein in the presence of C=O (**Figure S5**).

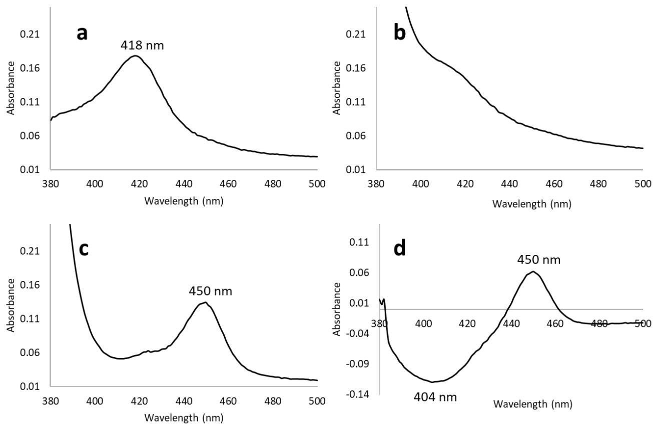


**Figure S5:** UV-Vis spectroscopic analysis of PtnJ. a: Purified PtnJ (25mM Tris-HCl, pH 8.0); b: Purified PtnJ + sodium dithionite; c: Purified PtnJ + CO and sodium dithionite; d: C=O difference spectrum.

##### Analysis of PtnJ-catalysed hydroxylation of filipin III

Filipin III was purchased from Sigma-Aldrich (≥ 85% purity) and dissolved in DMSO (10 mM final concentration) immediately prior to use. Purified PtnJ (6 µM) was incubated with NADPH (50 mM stock, 2 µL), ferredoxin (2 µL), ferredoxin reductase (2 µL) and freshly made filipin III solution (1 µL) in a total volume of 100 µL of Tris buffer (25 mM, pH 8). The reaction was incubated at room temperature for 3 h. 100 µL of methanol was added and the enzymatic mixture was allowed to stand for 5 min to precipitate the protein. After centrifugation at 13,200 rpm for 10 min, the supernatant was passed through a 0.2 µm filter and analysed by UHPLC-ESI-Q-TOF-MS, as described above.

**Table S1.** Summary of BGCs containing *ptnJ* orthologues

| Accession | Assembly ID | Species | Protein | % ID | Comments |
|---|---|---|---|---|---|
| NZ_FNTD01000004 | GCF_900104815 | *Streptomyces misionensis* DSM 40306 | WP_074995140.1 | 76.049 | PKS predicted to assemble same skeleton as *ptn*/*pen* cluster. Contains *ptnC*, *ptnD* and *ptnJ* orthologues. |
| NZ_FKJI03000020 | GCF_900079405 | *Streptomyces* sp. F-1 | WP_070027702.1 | 89.383 | Incomplete cluster with *ptnJ* orthologue adjacent to PKS gene. Genomic context similar to *S. misionensis* cluster. |
| NZ_MUNF01000001 | GCF_002154555 | *Streptomyces murinus* NRRL B-2286 | WP_086805776.1 | 99.753 | Incomplete cluster with *ptnJ* orthologue adjacent to PKS genes. |
| NZ_JOFU01000020[10] | GCF_000718315 | *Streptomyces griseofuscus* NRRL B-5429 | WP_037656760.1 | 99.259 | Incomplete cluster with *ptnJ* orthologue adjacent to PKS gene. Genomic context similar to *Streptomyces* sp. 816 cluster. |
| NZ_CP017248 | GCF_001735805 | *Streptomyces puniciscabiei* TW1S1 | WP_069777072.1 | 77.531 | PKS predicted to assemble same skeleton as *ptn*/*pen* cluster. Contains *ptnC*, *ptnD* and *ptnJ* orthologues. |
| NZ_LNBE01000002 | GCF_001445655 | *Streptomyces* sp. MBT76 | WP_058042043.1 | 88.642 | Incomplete cluster with *ptnJ* orthologue adjacent to likely misassembled PKS genes. |
| NZ_CP022438[11] | GCF_002777535 | *Streptomyces peucetius* subsp. *caesius* ATCC 27952 | WP_100109805.1 | 87.16 | PKS predicted to assemble same skeleton as *ptn*/*pen* cluster. Contains *ptnC* and *ptnJ* orthologues. |

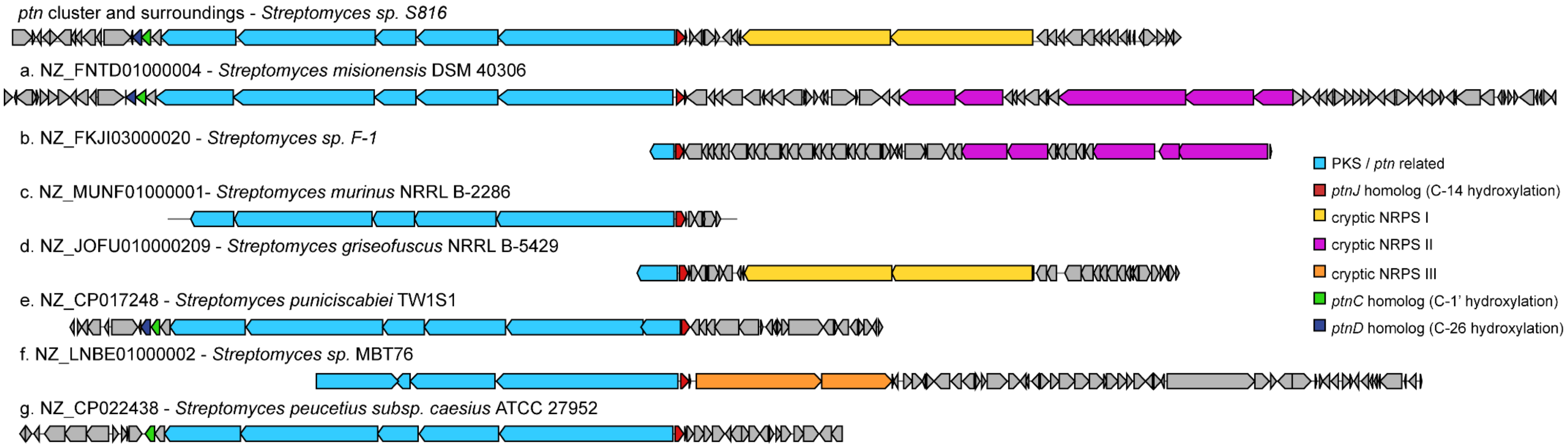


**Figure S6:** Comparison of gene clusters containing *ptnJ* orthologues in *Streptomyces* genomes.
